# A comprehensive prediction of transcript isoforms in 19 chicken tissues by Oxford Nanopore long-read sequencing

**DOI:** 10.1101/2022.07.18.500519

**Authors:** Dailu Guan, Michelle M. Halstead, Alma D. Islas-Trejo, Daniel E. Goszczynski, Hans H. Cheng, Pablo Ross, Huaijun Zhou

**Affiliations:** Department of Animal Science, University of California Davis, Davis, CA, 95616 USA; USDA, ARS, USNPRC, Avian Disease and Oncology Laboratory, East Lansing, MI 48823, USA

**Keywords:** transcriptome, annotation, transcript isoform, Nanopore, long-read sequencing, chicken

## Abstract

To comprehensively identify and annotate transcript isoforms in the chicken genome, we generated Nanopore long-read sequencing data from a diverse set of 19 chicken tissues comprising 68 samples collected from experimental line 6 × line 7 F_1_ adult males and females. More than 23.8 million reads with mean read length of 790 bases and average quality of 18.2 were generated. The annotation and subsequent filtering resulted in identification of 55,382 transcripts with mean length of 1,700 bases at 40,547 loci, representing ∼1.4 transcripts per locus. Among them, we predicted 30,967 potential coding transcripts at 19,461 loci and 16,495 potential lncRNA transcripts at 15,512 loci. Compared to reference annotations, we found 52% of annotated transcripts could partially to fully match while 47% were novel and potentially transcribed from lncRNA loci. Based on our annotation, we quantified transcript expression across tissues and found brain tissues (i.e. cerebellum, cortex) expressed highest number of transcripts and loci. The further tissue specificity revealed that ∼22% of the transcripts displaying tissue specificity. Of them, the reproductive tissues (i.e. testis, ovary) contained the most tissue-specific transcripts. Despite sequencing 68 transcriptomes derived from 19 tissues, still ∼20% of Ensembl reference loci were not detected. This suggests that including additional samples from different cell types, developmental and physiological conditions, is needed to fully annotate the chicken genome. The application of Nanopore sequencing transcriptomes in this study demonstrated the usefulness of long-read data in discovering additional novel loci (e.g., lncRNA loci) and resolving complex transcripts (e.g., the longest transcript for the *TTN* locus).

## 1 Introduction

Chicken (*Gallus gallus domesticus*) is one of the most widespread and common domesticated farm animals for egg and meat production, with a total population of 37.2 billion in stocks for the year 2020 (http://www.fao.org/). As the most popular studied bird species, moreover, its importance to the study of evolution, development, immunology, etc. is self-evident. In 2004, the first draft whole chicken genome was assembled with an estimated set of 20-23,000 protein-coding genes (PCGs) (Hillier et al., 2004). This effort offered a genome-wide view for understanding the configuration of the chicken genome, and the evolution of coding and noncoding vertebrate genomes. Since then, continuous efforts have been made to improve the completeness of chicken genome. For instance, Warren et al. (2017) added an additional 183 Mb sequences and assembled chromosomes 30-33 for the chicken reference genome. To fill the gaps of the chicken reference genome, recently two pangenomes were built that reported additional sequences absent from the GRCg6a reference genome (Wang et al., 2021a; Li et al., 2022).

The functional annotation of the chicken genome is also being produced in parallel. The two most commonly used databases, i.e. Ensembl (https://uswest.ensembl.org) and National Center for Biotechnology Information (NCBI, https://www.ncbi.nlm.nih.gov/) regularly update the chicken genome annotation. For instance, the Ensembl release (V102) included 16,779 PCGs and 39,288 transcripts, representing 2.34 transcripts per gene. Compared to the human ∼10 transcripts per gene, this estimate is quite low. The high estimate in human is partly attributed to the global efforts, such as GENCODE, which is part of the ENCODE (ENCyclopedia Of DNA Elements) consortium which aims to identify and classify all gene features in the human and mouse genomes. In farm animals, likewise, the consortium of the Functional Annotation of ANimal Genome (FAANG) was formed in order to improve the annotation of livestock genomes (Giuffra et al., 2019; Clark et al., 2020). In prior work, Kern et al. (2021) annotated noncoding genomes of three important livestock including chicken, and predicted 29,526 regulatory elements-gene interactions in chickens. In addition, Kern et al. (2018) also identified a total of 9,393 lncRNAs (including 5,288 novel lncRNAs) by utilizing short-read transcriptomes from eight chicken tissues.

The transcribed genomic region, though it only accounts for ∼3% of the genome, is very complex due to the alternative usage of transcription start, splicing, and polyadenylation sites. Alternative splicing has been shown to play important roles in evolution, phenotypic diversity, and organ development (Keren et al., 2010; Baralle and Giudice, 2017; Wright et al., 2022). For example, Yu et al. (2019) identified five alternative splicing variants of the *TYR* gene that were associated with skin melanogenesis in chickens. To annotate these features, transcriptome profiling provides important and useful resources (Yandell and Ence, 2012). For example, Jehl et al. (2020) annotated additional 1,199 PCGs and 13,009 long non-coding RNA genes (Compared to Ensembl V94) using 364 short-read transcriptomes derived from 25 chicken tissues. In human, a comprehensive annotation using transcriptomes of 41 tissues generated by Genotype-Tissue Expression (GTEx) Consortium improved transcript prediction for 13,429 genes, including 1,831 (63%) Online Mendelian Inheritance in Man (OMIM) genes and 317 neurodegeneration-associated genes (Zhang et al., 2020). This analysis demonstrated that a detailed annotation is better for understanding the phenome-to-genome connections. Although the short-read sequencing is widely used for annotating human and animal genomes, it is difficult to accurately resolve the complex structure of transcript isoforms. Chen et al. (2021), for instance, demonstrated that Nanopore long-read transcriptome sequencing classified individual isoforms better than Illumina short-reads despite they generated comparable gene expression estimates.

The contiguity of the long-read sequencing technology can sequence full-length transcript, thus is better suitable for dissecting the complexity of transcript structure compared to short-read sequencing (Muret et al., 2017). The Iso-Seq by the Pacific Biosciences is long-read sequencing technology that is widely used in profiling full-length transcriptome in human (Kuo et al., 2020), pig (Beiki et al., 2019), rabbit (Chen et al., 2017). In chickens, Thomas et al. (2014) used Iso-Seq long-read sequencing and identified 9,221 new transcript isoforms in embryonic chicken heart tissue. Later on, Kuo et al. (2017) annotated 64,277 additional distinct transcripts (55,315 in brain and 9,206 in embryo) using Iso-Seq plus 5′ cap selection in chicken brain and embryo tissues. However, the few tissues studied (only brain and embryo included in Kuo et al. (2017)) make it difficult to capture the diversity of chicken transcript variations.

The Oxford Nanopore Technologies has provided an alternative long-read sequencing approach (Amarasinghe et al., 2020), which has been applied in cattle (Halstead et al., 2021), duck (Lin et al., 2021) and many other species, but not yet in chickens. The Nanopore long-read sequencing allows for accurate identification and quantification of transcript isoforms and for resolving complex isoforms (Byrne et al., 2017; Soneson et al., 2019; Chen et al., 2021). In this study, we aimed to identify and characterize transcripts in a diverse set of chicken tissues, including cerebellum, hypothalamus, cortex, duodenum, jejunum, ileum, cecum, colon, testis, ovary, adipose, gizzard, heart, kidney, liver, lung, muscle, spleen and thymus, using Oxford Nanopore long-read sequencing technology. The data generated from this study will be a valuable source to improve our understanding of the complexity of the chicken transcriptome, and also aid in dissecting the connection of gene expression and phenotypic traits.

## 2 Methods and Materials

### 2.1 Sample collection

All animals and samples used in this study were obtained in concordance with the Protocol for Animal Care and Use no. 18464 (approved by Institutional Animal Care and Use Committee at the University of California at Davis). All tissues were from one of two FAANG pilot projects (FarmENCODE) (Tixier-Boichard et al., 2021). In brief, ADOL experimental White Leghorn lines 6_3_ and 7_2_ were intermated to produce F_1_ progeny, and 4 male and 2 female individuals were euthanized at 20 weeks of age. Tissues were collected within 1-2 hours and stored at -80 °C until further use.

### 2.2 RNA extraction and library preparation

RNA extraction and library preparation were done by following the protocols reported in (Halstead et al., 2021). Briefly, frozen tissues were mashed using a pestle in a mortar filled with liquid nitrogen. Then, Trizol reagent (Invitrogen, Carlsbad, CA, United States) was added to extract total RNA using the Direct-zol RNA Mini Prep Plus kit (Zymo Research, Irvine, CA, United States). The integrity and quality of extracted RNA were checked using an Experion electrophoresis system (Bio-Rad, Hercules, CA, United States) and those passing quality control were used for library preparation. First, 50 ng of total RNA in a volume of 9 μl was mixed with 1 μl 10 μM VNP primer, 1 μl 10 mM dNTPs for incubation 5 min at 65 °C. The products were used for strand-switching and reverse transcription reactions. Then, barcodes were ligated to the cDNA products generated from the last step using the Oxford Nanopore PCR barcoding expansion 1-96 kit (cat. no. EXP-PBC096), which were further ligated with adapters from the SQK-DCS109 kit following the manufacturer’s guidelines. Products were loaded onto a PromethION flow cell (vR9.4.1) for sequencing.

### 2.3 Base calling, quality control and preprocessing

After base calling and de-multiplexing with the ont-guppy-for-minknow (v3.0.5) tool (https://nanoporetech.com/), we run quality control using NanoPlot (v1.0.0) software in order to summarize read length, average quality, among others. Then, the Pychopper software (https://github.com/nanoporetech/pychopper) was employed to identify and orient full-length reads, which were mapped against the reference genomes (GRCg6a, Ensembl V102) with options of “-ax splice -uf -k14 -G 1000000” using the minimap2 software (Li, 2018). We discarded reads with a minimum quality score of 10 using SAMtools (v1.9) (Li et al., 2009) and counted the gene expression using the HTSeq 0.13.5 software (Anders et al., 2015). Read counts were then normalized into variance stabilizing transformation (VST), which was used for sample clustering analysis with the function of “plotPCA” implemented in the DEseq2 software (Love et al., 2014).

### 2.4 Reference-guided prediction of transcript isoforms

To predict transcripts, we used a computational pipeline supported by the Oxford Nanopore Technology community (https://github.com/nanoporetech/pipeline-nanopore-ref-isoforms). Briefly, the oriented full-length reads with fastq format were pooled together and then mapped against the Ensembl annotation (GRCg6a, V102) using minimap2 (Li, 2018) in order to carry out a reference-guided transcriptome assembly. The mapped reads were then used to annotate transcripts using the StringTie2 software (Kovaka et al., 2019) in the long-read mode (with the option of “-L”). Transcripts on unplaced scaffolds, as well as those with exon coverage < 100% and read depth < 2 were excluded. Then we only keep single-exon transcripts with expression TPM > 1 in > 2 samples of a tissue, and multi-exon transcripts with expression TPM >0.1 in >2 samples of a tissue. After that, we further excluded transcripts categorized as potential artifacts (see **Comparing predicted transcripts to previous annotations** section).

### 2.5 Prediction of coding and non-coding transcripts and loci

To predict the coding potential of predicted transcripts, we employed the TransDecoder (https://github.com/TransDecoder/TransDecoder) and CPP2 software (Kang et al., 2017). The final predictions are composed of either TransDecoder or CPP2 ones. After prediction of coding potential, we obtained the list of non-coding transcripts, which were further used for predicting whether they are lncRNA loci using the FEElnc software (Wucher et al., 2017).

### 2.6 Comparing predicted transcripts to previous annotations

The predicted transcripts were compared to the Ensembl (V102) and NCBI reference (V105) annotations using GffCompare tool (version 0.11) (Pertea and Pertea, 2020) and classified into 14 classes. According to Halstead et al. (2021), the predicted transcripts classified into four categories: exact match (class code “=”), which means the intron chains of our annotated transcripts can exactly match to reference annotations; novel isoform (class codes ‘c,’ ‘k,’ ‘j,’ ‘m,’ ‘n,’ or ‘o’), which means predicted transcript can not match a reference transcript but can match a reference gene; novel loci (class codes ‘i,’ ‘u,’ ‘y,’ or ‘x’), which means predicted transcript can not match either a reference transcript or a reference locus; and potential artifacts (class codes ‘e,’ ‘s,’ or ‘p’), which are possibly due to mapping error, e.g. pre-mRNA fragments, polymerase run-on, etc. To compare our prediction with novel transcripts reported by Thomas et al. (2014), we first converted positions of their transcripts from galGal4 to GRCg6a using the liftover software (Kuhn et al., 2013). Then the GffCompare tool was used for comparing our annotation to their transcripts.

### 2.7 Quantification of predicted transcripts

We extracted sequences of predicted transcripts using GffRead v0.12 (Pertea and Pertea, 2020), which constituted a reference transcriptome in the FASTA format. Then, we mapped the full-length reads generated by Pychopper (https://github.com/nanoporetech/pychopper) to the predicted transcriptome using minimap2 (v2.1) (Li, 2018). The transcript expression was quantified using Nanocount (v0.2.4) (Leger, 2020). Based on the metric of the transcripts per million (TPM), we categorized transcripts as highly (average TPM ≥ 10), moderately (1 ≤ average TPM < 10), and lowly expressed (average TPM < 1) (Halstead et al., 2021).

### 2.8 Tissue-specificity analysis

The tissue specificity of transcripts expression across tissues were evaluated by using a tissue specificity index (*TSI*) (Julien et al., 2012; Halstead et al., 2021):

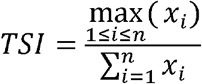

Where *x*_*i*_ is an average of transcript expression (TPM) in a given tissue, n is the number of tissues. Transcripts were then categorized as tissue-specific (TSI ≥ 0.8), broadly expressed (TSI < 0.5), or biased towards a group of tissues (0.5 ≤ TSI < 0.8). To reveal functional biology of tissue specific transcripts, we extracted tissue-specific transcript sequences and blast them against the SwissProt (protein sequence database, V5) using the Diamond blastx tool (v2.0.11.149) (Buchfink et al., 2015). We then carried out functional enrichment (only considering Gene Ontology Biological Process terms) using the matched UniProt identifiers using the PANTHER tool (Mi et al., 2013). The false discovery rate (FDR) approach (Benjamini and Hochberg, 1995) was used for multiple testing corrections and FDR value less than 0.05 was set as the significance threshold.

### 2.9 Differential alternative splicing analysis

To detect differential alternative splicing (DAS) events, we employed the LIQA software (Hu et al., 2021). Based on our annotation, we quantified isoform expression using the “quantify” function. Then the DAS events between tissues were detected using the “diff” within the LIQA tool (Hu et al., 2021). After multiple testing correction using FDR approach (Benjamini and Hochberg, 1995), the threshold of significance was set as FDR < 0.05.

## 3 Results

To comprehensively annotate transcripts of the chicken genome, we sequenced 68 samples comprising of 19 tissues collected from six individuals (two females: CC and CD; and four males: CA, CB, M1, M2) (**Supplementary Table 1**). The tissues collected were cerebellum, hypothalamus, cortex, duodenum, jejunum, ileum, cecum, colon, testis, ovary, adipose, gizzard, heart, kidney, liver, lung, muscle, spleen and thymus. Sequencing generated a total of 23.8 million reads, with an average of 344,650 reads per tissue and an average length of 790 bp (**Figure 1a, Supplementary Table 2**).

**Figure 1.**
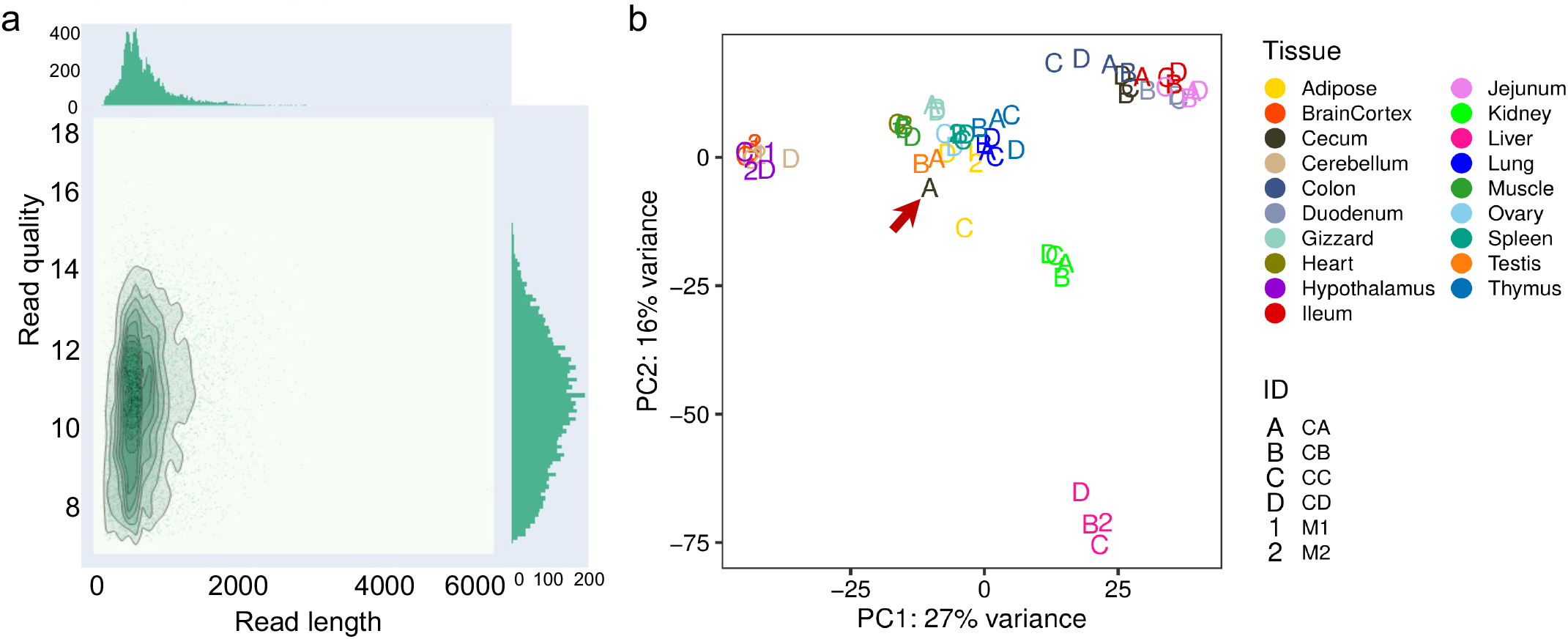
(a) Bivariate plot (De Coster et al., 2018) depicting read length (x-axis) and quality (y-axis) of Nanopore long-read sequencing in 68 samples (b) Principal component analysis of 68 chicken Nanopore long-read transcriptomes. The red arrow indicated the sample, CA_Cecum, which was not clustered with other samples from the cecum tissue.

Principal component analysis (PCA) and hierarchical clustering of mapped sequencing reads to the Ensembl annotation (GRCg6a, version 102) revealed that samples generally clustered according to the origin of tissues or organs as expected (**Figure 1b** and **Supplementary Figure 1**). Moreover, we found samples from the same biological system tend to cluster together, such as brain cortex, cerebellum and hypothalamus from the central neural system; cecum, colon, duodenum, ileum, and jejunum from the intestinal system (**Figure 1b**). However, an outlier (i.e., Cecum_CA) in the PCA plot and hierarchical clustering (indicated by a red arrow in **Figure 1b** and **Supplementary Figure 2**) was observed, even separated from the same cecum tissue. The summary statistic indicates that the unexpected clustering is possibly due to the insufficient sequencing depth (number of reads = 1,279), which was lower than the rest (average number of reads for remaining samples = 355,008, **Supplementary Figure 2**). However, 895 out of 1,279 reads from Cecum_CA reads aligned to the GRCg6a genome, corresponding to a mapping rate of 70%. In the light of these analyses, we included the Cecum_CA sample in transcript prediction, but not in the transcript expression analysis, e.g., tissue specificity of transcript expression, differential alternative splicing (DAS) analysis.

To assemble potential transcripts, we identified, oriented and trimmed full-length reads using the Pychopper v2 software. Then, the StringTie tool with the long read mode was used for predicting transcripts (https://github.com/nanoporetech/pipeline-nanopore-ref-isoforms). As a result, 79,757 transcripts in 54,551 loci in total were identified. After filtering out transcripts on unplaced scaffolds, as well as those with exon coverage < 100% and read depth < 2, we obtained 74,665 transcripts in 50,569 loci, of which there were 45,132 multi-exon and 29,533 single-exon transcripts. Moreover, we required multi-exon transcripts with expression TPM (Transcripts Per Million) > 0.1 and single-exon transcripts with expression TPM > 1 in at least 2 sample of a tissue. By doing so, there were 61,556 transcripts in 45,284 loci remained. To further exclude potential artifacts, we compared assembled transcripts with NCBI (V105) and Ensembl (V102) reference annotations. The result is shown in **Figure 2a** and **Table 1** (see **Methods**). Overall, we found ∼14% of predicted transcripts can exactly match the reference annotations (**Figure 2a**). With the Ensembl annotation, 77% of them were considered as novel transcripts, either novel isoforms (35%) or novel loci (42%). In addition, ∼8% were potential artifacts, which possibly caused by pre-mRNA fragment, polymerase run-on or mapping error (**Figure 2a**). After excluding these potential artifacts, we finally kept 55,382 transcripts in 40,547 loci, representing 1.4 transcripts per locus (**Supplementary Data 1**).

**Figure 2.**
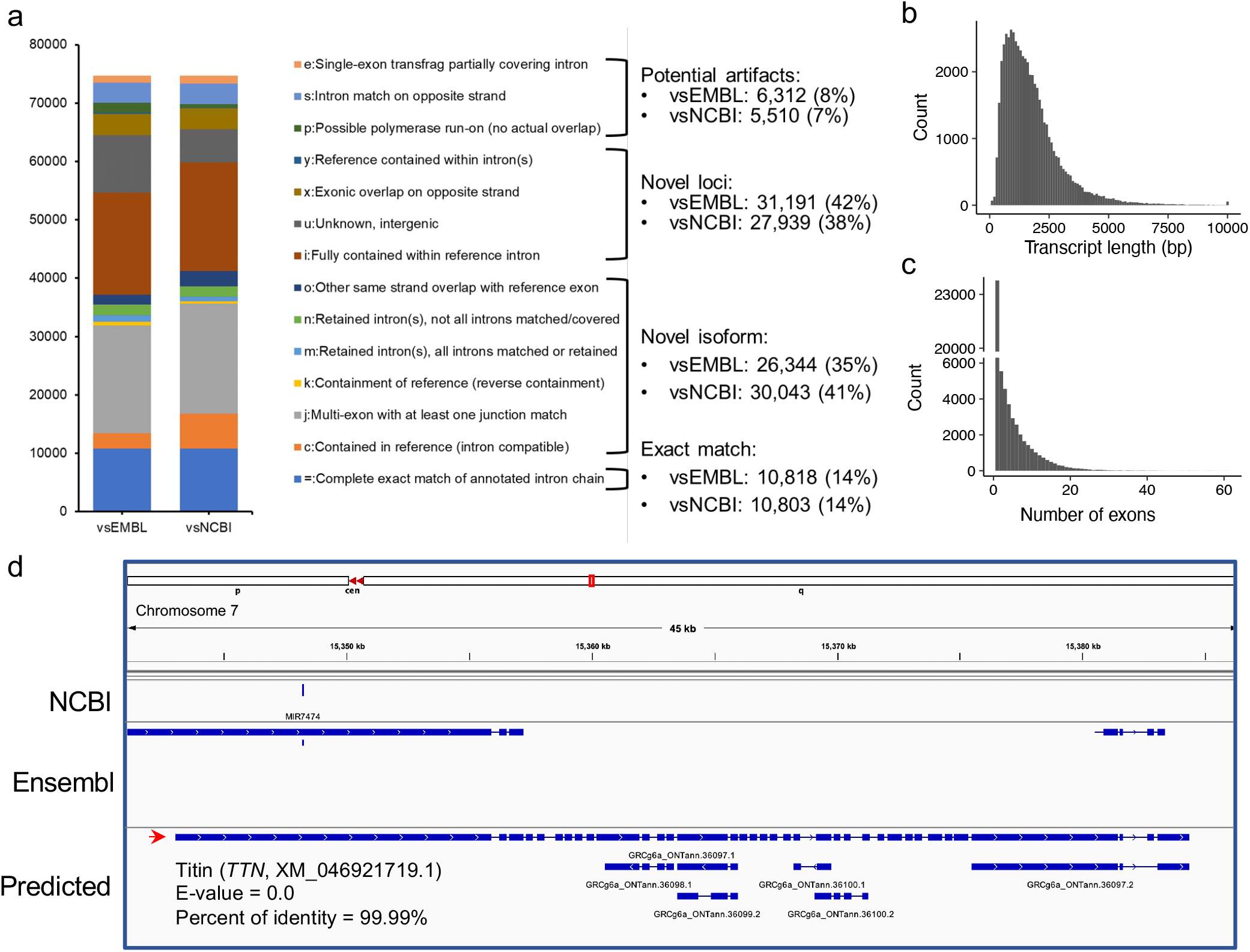
(a) Comparisons of predicted transcripts against Ensembl (V102, vsEMBL) and NCBI annotation (V105, vsNCBI). The transcripts were classified according to the GffCompare software (Pertea and Pertea, 2020). The panels (b) to (c) depict the distributions of predicted transcript length and exon numbers, respectively. (d) A screenshot showing the predicted longest transcript, which is located on chromosome 7 (15,343,033-15,384,347). Blast analysis indicated the transcript matched to the *TTN* gene locus encoding the titin protein.

**Table 1.**
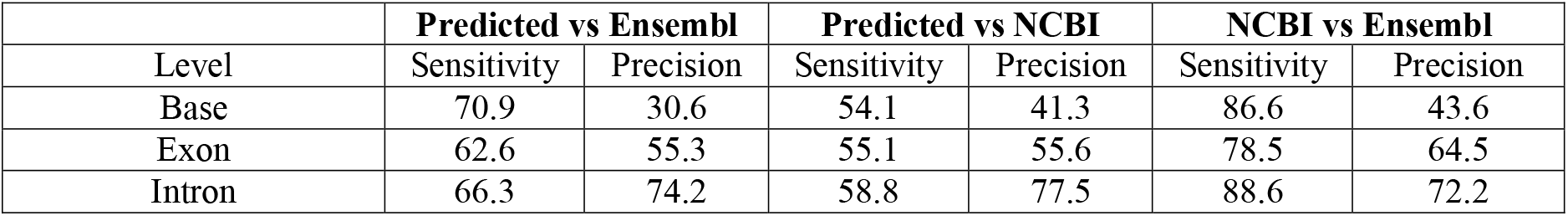
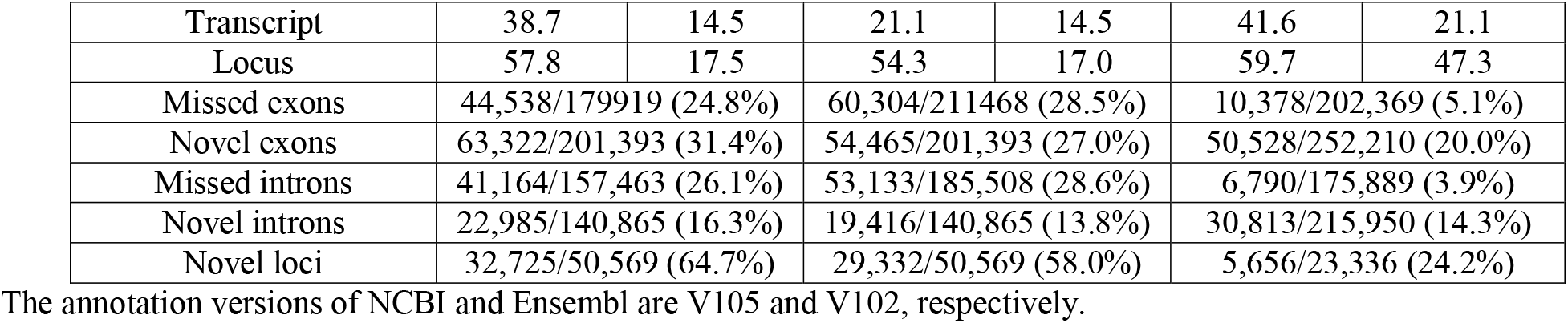
Comparison of reference and predicted transcripts using GffCompare tool

The length of predicted transcripts ranged from 49 to 34,500 bp, with a mean length of 1,767 bp (**Figure 2b**). The longest transcript, for instance, is located on chromosome 7 (15,343,033-15,384,347), which highly matched to the *TTN* gene encoding the giant protein titin (NCBI reference sequence XM_046921719.1, E-value = 0.0, percent of identity = 99.99%) (**Figure 2d**). This protein plays important roles in the movement of skeletal muscle, but its gene locus has not been annotated in both NCBI (V105) and Ensembl (V102) GRCg6a reference annotations (**Figure 2d**). Moreover, we found the annotated 55,382 transcripts are supported by 171,651 unique exons, with an average estimate of 4.34 exons per transcript (**Figure 2c**).

To predict the coding potential of predicted transcripts, we employed the CPC2 and TransDecoder software. The former predicted 21,984 transcripts at 12,999 loci with coding potential, and the latter one predicted open reading frames for 30,727 transcripts corresponding to 19,306 loci. In total, we predicted 30,967 uniquely potential coding transcripts at 19,461 loci, representing 1.6 transcripts per locus (**Supplementary Table 3**). Furthermore, we surveyed whether the remaining 24,415 transcripts were long non-coding RNAs (lncRNAs). To do so, we employed the FEELnc software and found 16,495 potential lncRNA transcripts at 15,512 loci (**Supplementary Table 3**).

We compared our prediction to two reference annotations and found the number of transcripts per locus of our annotation (∼1.4) is lower than reference annotations (Ensembl v102: ∼1.8 transcripts per locus; NCBI v105: ∼3.3 transcripts per locus), but we predicted ∼20K more loci, of which a substantial proportion is lncRNA loci (**Figures 3a** and **3c**). At the transcript level, we classified transcripts into three categories (see **Methods**): 1) exact match: predicted transcripts completely match to reference annotations; 2) novel isoform: predicted transcripts do not match reference transcripts but match reference loci; 3) novel loci: predicted transcripts do not match any reference loci and transcripts (**Figure 3b**). Concordantly, we found our prediction identified high proportion of “novel loci” transcripts (47%), followed by “novel isoforms” (37%) when comparing to Ensembl annotation (V102) (**Figure 3b**). A similar pattern was observed when comparing to NCBI annotation (**Supplementary Figure 3**). By further comparing lncRNA loci predicted in this study with those predicted by Jehl et al. (2020), we found ∼ 83% of our predicted lncRNA transcripts can match their annotations (**Supplementary Figure 4**). Thomas et al. (2014) also reported 9K novel transcripts from long-read sequenced embryonic chicken heart transcriptomes. By comparing these available novel transcripts to our annotation, we found 89% of them can completely or partially match to our annotation, while there were still 1,000 transcripts categorized as “novel loci” (**Supplementary Figure 5**). Moreover, we found the transcripts grouped into the “novel isoform” and “novel loci” categories tend to be lowly expressed, while the expressions of transcripts in “exact match” group are higher (**Figure 3d**).

**Figure 3.**
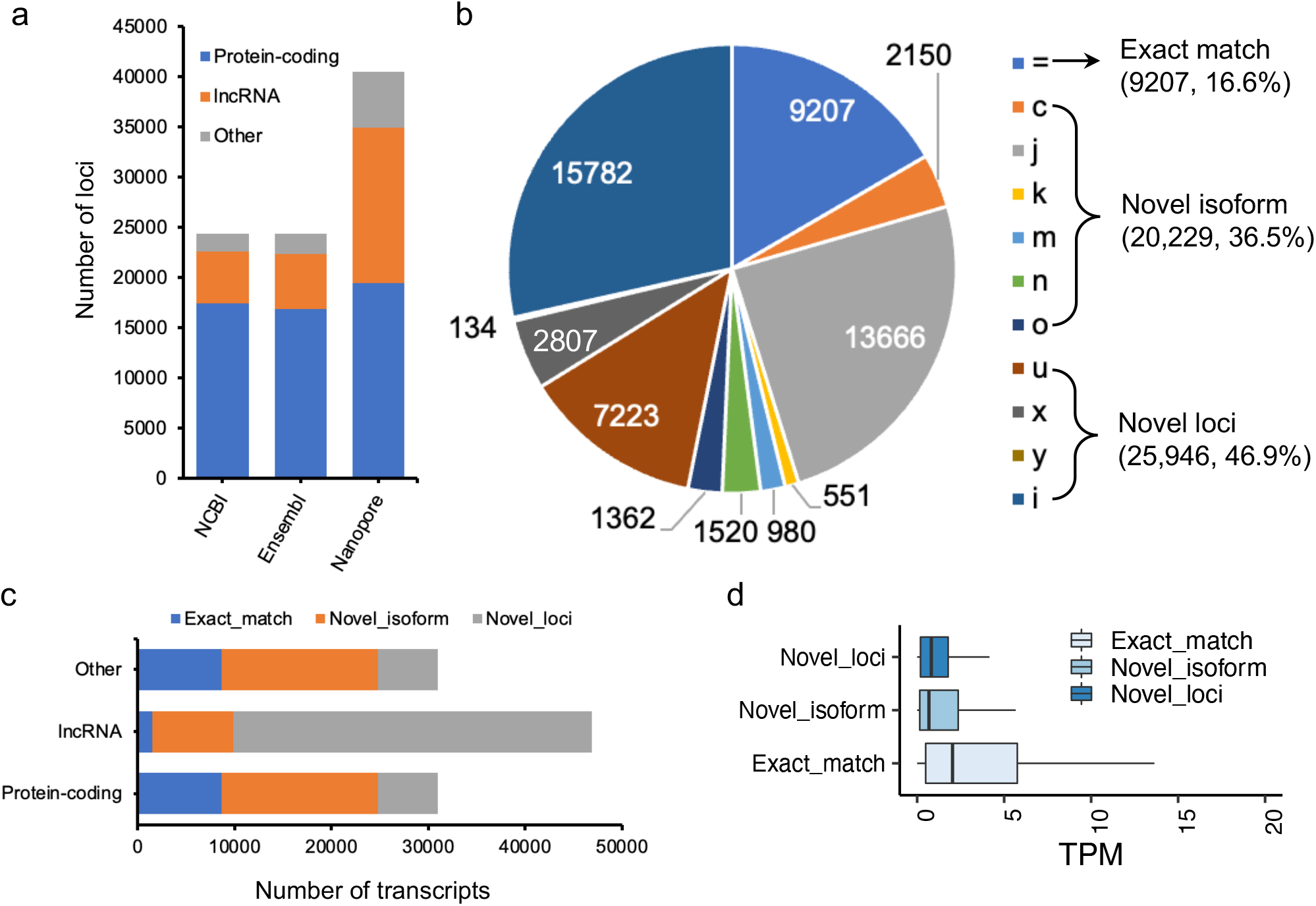
(a) Number of loci in NCBI (V105), Ensembl (V102) and our annotations. (b) Pie chart depicting GffCompare types to Ensembl annotation (V102). See **Methods** for explanation of the type codes. (c) Number of transcripts as a function of protein-coding, lncRNA, and other non-coding loci. (d) Transcript expression measured as transcript per million (TPM) as a function of different types of transcripts classified by GffCompare (See **Methods**).

Considering the largest set of tissues used, we then sought to identify tissue-specifically expressed transcripts. By quantifying transcript expressions, we found the number of expressed transcripts and loci ranged from 14,841 (liver) to 28,648 (cerebellum), and from 10,285 (liver) to 21,662 (cerebellum), respectively (**Supplementary Figure 6**). The tissue specificity index (TSI) indicated that the set of “exact match” transcripts tend to be lowly tissue-specific, while “novel isoform” and “novel loci” transcripts are highly tissue-specific (**Figure 4a**). We observed that the set of transcripts with low expression tended to have high tissue-specificity, while in contrast, highly expressed transcripts are commonly found across tissues (**Figure 4b**). Moreover, we identified tissue-specific transcripts and found the reproductive tissues (i.e., testis and ovary) have high proportion of tissue-specific transcripts, followed by brain-related tissues (i.e., cerebellum and cortex) (**Figure 4c**). For instance, we identified a novel transcript located on chromosome 4 (52,482,563-52,492,561), which is specifically expressed in testis samples (**Figures 4d** and **4e**). This transcript was predicted as a sense intergenic lncRNA by using the FEELnc software (Wucher et al., 2017) (**Supplementary Tables 3** and **4**). To reveal the function of tissue-specific transcripts, we aligned sequences of tissue-specific transcripts to SwissProt (V5) database with the blastx function implemented in the Diamond tool (v2.0.11.149) (Buchfink et al., 2015). Then, the matched UniProt identifiers were used for carrying out functional enrichment analysis with the PANTHER tool (Mi et al., 2013). This analysis revealed that tissue-specific transcripts recapitulated the tissue biology (**Figure 5a, Supplementary Table 5**), such as muscle contraction, muscle cell differentiation enriched in muscle and heart tissues, trans-synaptic signaling and nervous system development in cerebellum and brain cortex, and B cell receptor signaling pathway in spleen (**Figure 5a, Supplementary Table 5**), a finding concordant with previous results (Yang et al., 2018; Fang et al., 2020).

**Figure 4.**
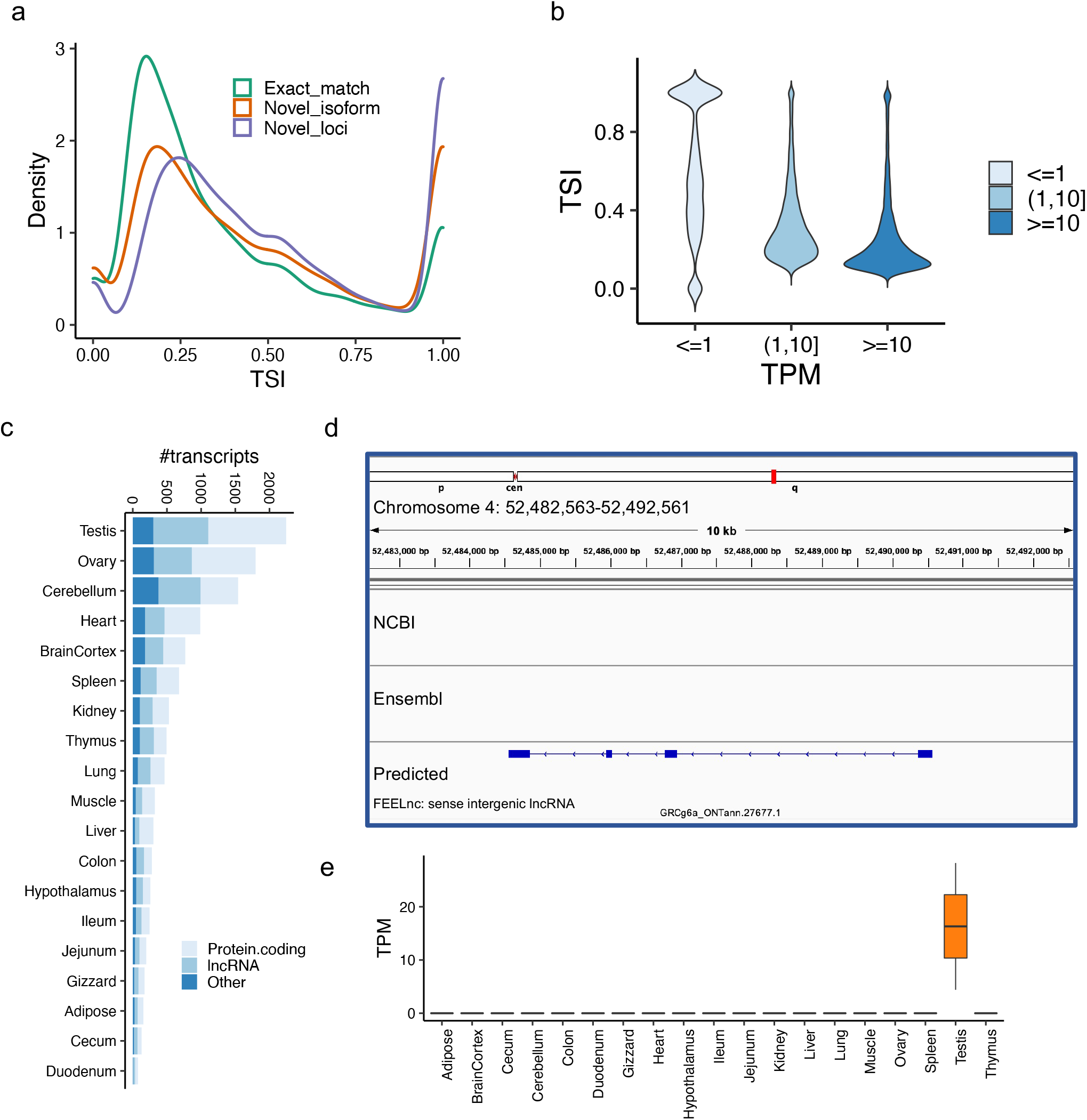
(a) Tissue specificity index (TSI) as a function of different types of transcripts classified by GffCompare (See **Methods**). (b) Transcript expression measured as transcript per million (TPM) as a function of tissue specificity index (TSI). We grouped transcripts according to their expressions (see **Methods**). (c) Number of tissue-specific transcripts in each tissue. (d) A screenshot showing a novel transcript only predicted by our data, which is located on chromosome 4 (52,482,563-52,492,561). The transcript is highly expressed in testis samples, but not any other tissue samples. The FEELnc predicted it as a sense intergenic lncRNA.

**Figure 5.**
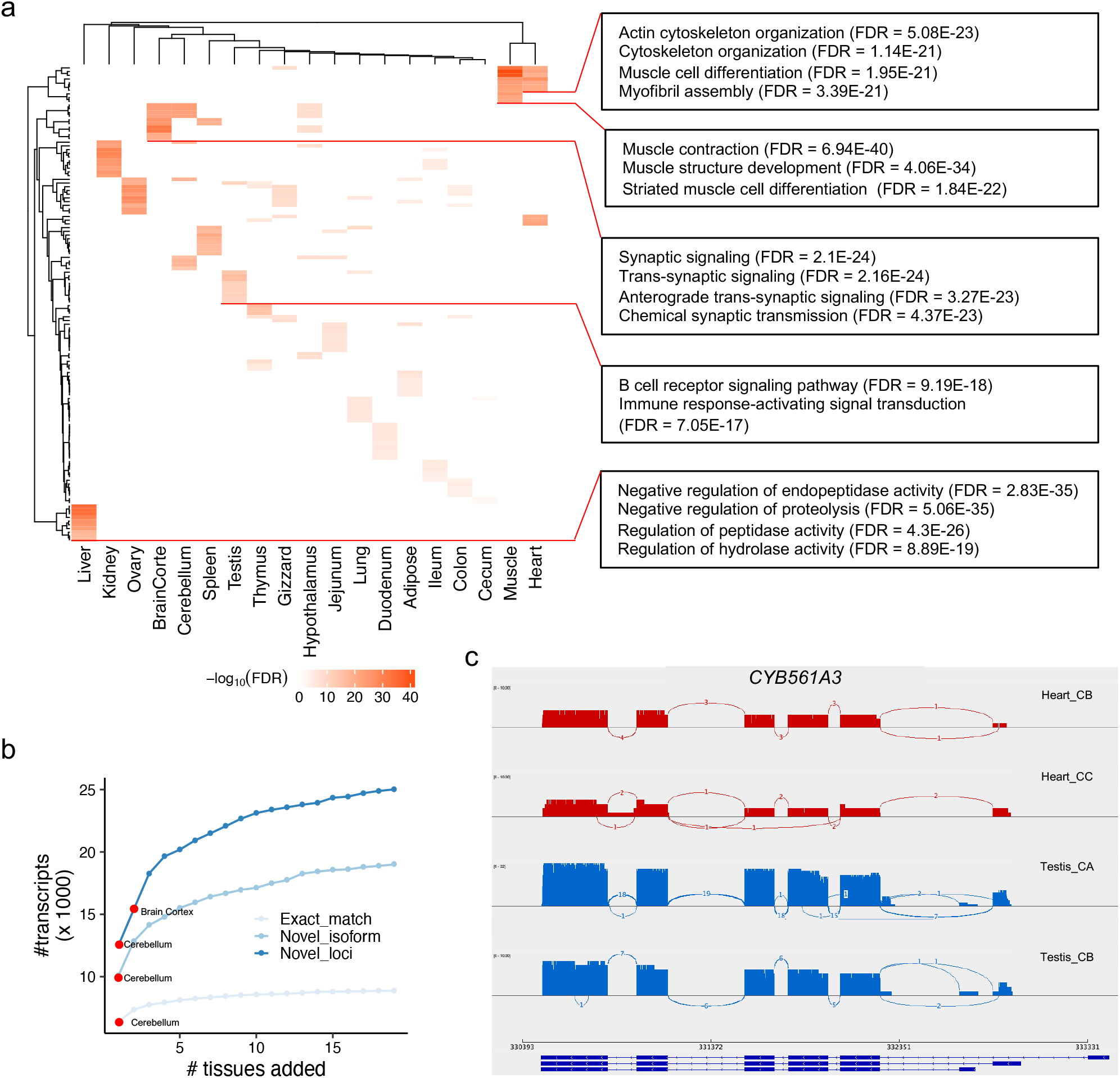
(a) Heatmap depicting the negative log_10_ FDR (false discovery rate) values for the top 10 Gene Ontology (GO) Biological Process terms. At the right side, we show several examples of GO terms, as well as their FDR values. (b) Number of unique transcripts detected as a function of tissues added. Transcripts are categories into three types (see **Methods**). (c). Sashimi plots of *CYB561A3* gene which showed DAS between heart (red) and testis (blue).

The utilization of large scale of tissues allows us to investigate which tissue is better to capture more transcripts and to annotate chicken genome. Herein we tried to detect the number of unique transcripts expressed as a function of more tissues added. By doing so, we found brain-related tissues (i.e., cerebellum and cortex) could detect the higher number of transcripts as expected (**Figure 5b, Supplementary Table 6**). In addition, our design including a diverse set of 19 chicken tissues offers the opportunity to analyze DAS events between chicken tissues. To do so, we quantified isoform expression and identified differential alternative splicing events using the LIQA software (Hu et al., 2021). The results are shown in **Supplementary Figure 7** and **Supplementary Table 7**. In total, we found a list of 4,211 loci showing DAS events between tissues (FDR < 0.05). For instance, the top significant locus is the *CYB561A3* gene showing DAS between heart and testis (FDR = 9.12E-16, **Figure 5c**). This gene encodes cytochrome B561 family member A3 whose functions are related cellular iron ion homeostasis and mitochondrial respiration (Wang et al., 2021b).

## 4 Discussion

A well-annotated chicken genome is essential in associating genomic variation to phenotypic variation, and there are a number of ongoing efforts through the Functional Annotation of Animal Genomes (FAANG) consortium (Andersson et al., 2015), primarily focus on non-coding functional elements in farm animals including chicken (Kern et al., 2021). In this study, using Oxford Nanopore long-read sequencing in 19 chicken tissues, we preliminary annotated 79,757 transcripts in 54,551 loci, while the subsequent filtering resulted in the exclusion of ∼2K transcripts. Finally, our prediction resulted in the identification of 55,382 clean transcripts derived from 40,547 loci, representing ∼1.4 transcripts per locus, an estimate lower than the Ensembl (∼1.8 transcripts per locus), and the NCBI annotations (∼3.3 transcripts per locus). The lower estimate in our study might be due to the higher number of annotated loci (N = 40,547), i.e. around 2.6-fold higher than both reference annotations.

The number of transcripts of loci predicted in this study is substantially higher than two reference annotations (Esembl V102: 27,955 transcripts in 15,305 loci; NCBI V105: 51,222 in 15,706 loci), while our prediction is lower than Kuo et al. who annotated 60K transcripts and 29K genes using the Iso-Seq approach (Kuo et al., 2017). Unfortunately, the unavailability of their annotation hinders us to exclusively make a comparison. Specifically, we predicted higher proportion of lncRNA loci, indicating that reference annotations are not annotated lncRNAs well. Indeed, Jehl et al. (2020) annotated additional 13,009 lncRNA genes (compared to Ensembl V94) using 364 chicken short-read transcriptomes derived from 25 tissues. Indeed, when we compared our lncRNA transcripts to Jehl et al. (2020), we found over 80% of them completely or partially match to their lncRNA loci. Still, our annotation contains 4,953 additional novel lncRNA transcripts despite we used the sample lncRNA prediction tool (FEELnc, Wucher et al., 2017), which may be due to the increased sensitivity of long-read sequencing (Lagarde et al., 2017). Moreover, we found > 89% of novel transcripts reported by Thomas et al. (2014) could match our prediction. These evidences collectively indicate our annotation is reliable.

Comparing to the reference annotations, we observed a higher percentage of novel loci (∼47%) than that of cattle (6% predicted transcripts did not match to a reference gene), whilst the exact matched transcripts predicted in this study were also lower (16% in our study vs. 21% in cattle), though the cattle study included more tissues (Halstead et al., 2021). Potential reasons are low number of samples, possible degradations of RNA samples or low sequencing depth. We also cannot rule out the possibility that the annotation of the bovine reference genome is better than the chicken one in the database. It should be noted that a substantial proportion of novel loci predicted by us are lncRNA loci which can to some extend match to a previous study (Jehl et al., 2020). These results suggest more efforts for annotating the chicken genome is needed in the future. The human genome is considered to be better-annotated than farm animals’, while 36.4% of full-length transcripts identified by long-read Iso-Seq methods are classified as “novel” in human cortex tissue (Leung et al., 2021). Using the same approach, another study also reported 17 to 55% of novel isoforms in human breast cancer samples (Veiga et al., 2022). These studies, together with ours, indicate long-read sequencing is better approach for discovering novel isoforms and being able to better annotate animal genomes.

The number of transcripts reported by this study, reference genome annotations, as well as by Kuo et al. (2017) varies widely, ranging from 27,955 to 74,665. Although sequencing depth could be one of reasons, another possible interpretation is that the number of detectable transcripts is tissue-dependent. Indeed, our study with similar sequencing depth also detected variable number of expressed transcripts across tissues, ranging from 14,841 (liver) to 28,648 (cerebellum). These observations suggest that including as diverse and many tissues as possible can detect tissue-specific transcripts and better annotate the chicken genome. It is reported that brain tissues have a higher level of alternative splicing, such as skipped exons, alternative 3’ splice site exons or 5’ splice site exons, (Yeo et al., 2004; Melé et al., 2015). Our analysis supported this notion, suggesting brain-related tissues are better for annotating an animal genome if available tissues are limited. The consistent pattern of the higher number of transcripts observed in brain possibly reflects the complexity of tissue biology (Naumova et al., 2013; Fang et al., 2020). Moreover, the whole embryo was also expected to include as many transcripts as possible since it contains all organs. Unfortunately, our study design did not include the whole embryo, but in Kuo et al. study they identified 55,932 transcripts in brain while only 9,368 transcripts in embryo (Kuo et al., 2017).

Although our study has annotated a substantial proportion of novel transcripts, there were still some limitations, e.g., our study only includes a single developmental stage (adult). Previous reports indicate that detecting gene expression using long-read sequencing approaches requires lower number of reads, such as Nanopore sequencing needs ∼ 40-fold less reads or ∼ 8-fold less bases than Illumina technology, which required over 36 million reads for accurately quantifying highly expressed genes (FPKM > 10), and over 80 million reads for lowly expressed genes (FPKM < 10) (Sims et al., 2014; Su et al., 2014; Oikonomopoulos et al., 2020). Based on that estimate, at least 7.5 million long-reads are likely to be required per tissue, while this will be the cost-prohibitive given the cost of Nanopore long-read sequencing we did in 2019 with so many samples. This indicates our study, very possibly, missed a proportion of transcripts due to the low sequencing depth, though we reported a higher number of transcripts and loci than reference annotations. This is also reflected by the ratio that each gene can only produce 1.4 transcripts per locus based on our data, while each human gene can produce ∼10 splicing transcripts (Mathur et al., 2019). With continued DNA technologies development in cost-effective, tissues from more developmental stages and physiological status, and more in-depth sequencing on full-length transcriptome are warranted to improve the annotation of transcript isoforms in the chicken genome.

## 1 Conflict of Interest

None.

## 2 Author Contributions

HZ and PR conceived and designed the experiments. ADI, DEG, HC collected samples and carried out nanopore sequencing experiments. DG and MM developed the computational pipeline and analyzed all data. DG and HJ wrote the paper. All authors read, edited and approved the final manuscript.

## 3 Funding

Funding for sample collection was provided by the United States Department of Agriculture, National Institute of Food and Agriculture, Agriculture and Food Research Initiative grant no. 2015-43567015-22940 awarded to HZ and PR. Funding for library generation, and sequencing, was provided by the United States Department of Agriculture, National Institute of Food and Agriculture, Agriculture and Food Research Initiative grant no. 2017-67015-26297 awarded to PR and HZ. Funding for bioinformatics analysis was provided by Agriculture and Food Research Initiative grant no. 2020-67015-31175 awarded to HZ.

## 4 Acknowledgments

We thank Dr. Ying Wang, Dr. Perot Saelao, and Ganrea Chanthavixay for their assistance in sample collection, organization, and storage. We also thank the staff at the University of California, Davis, DNA Technologies Core for their guidance and assistance with Nanopore library construction and sequencing.

## 6 Supplementary Material

**Supplementary data 1** Predicted transcripts in the General Feature Format (GTF) format

**Supplementary Table 1** Information about tissue sampling used in this study

**Supplementary Table 2** Summary statistics of sequencing samples

**Supplementary Table 3** Predicted transcript types (including protein-coding, lncRNA and other non-coding)

**Supplementary Table 4** A list of tissue-specific transcripts

**Supplementary Table 5** Functional enrichment of tissue-specific transcripts (only Biological Process of Gene Ontology terms)

**Supplementary Table 6** Number of unique transcripts detected when adding more tissues

**Supplementary Table 7** A list of loci showing differential alternative splicing (DAS) events between tissues

**Supplementary Figure 1** Hierarchical clustering of samples used in this study. The dendrogram is built based on gene expressions quantified with Transcripts Per Million (TPM > 0.1). The distance between individuals is indicated by 1-r, where r is the Pearson correlation coefficient

**Supplementary Figure 2** Dotplot depicting the number of sequencing reads (x-axis) and the number of expressed genes (y-axis). A given gene was considered as expressed when Transcripts Per Million (TPM) > 0.1. The red text indicates the outlier sample in principal component analysis (PCA) plot (Figures 1b) and hierarchical clustering (Supplementary Figure 1).

**Supplementary Figure 3** GffCompare types when comparing our predicted transcripts to NCBI annotation (V105)

**Supplementary Figure 4** GffCompare types when comparing protein-coding (a) and lncRNA loci (b) predicted in this study with those predicted in Jehl et al., (2020).

**Supplementary Figure 5** GffCompare types when comparing novel transcripts reported by Thomas et al. (2014) to our annotation.

**Supplementary Figure 6** Number of expression loci and transcripts (TPM > 0.1) across tissues

**Supplementary Figure 7** Number of loci showing differential alternative splicing (DAS) between tissues

## 7 Data Availability Statement

The Nanopore sequencing data are accessible in the Sequence Read Archive (SRA) database of the National Center for Biotechnology Information with the identifier PRJNA671673 (https://www.ncbi.nlm.nih.gov/bioproject/PRJNA671673). The code used for annotating full-length transcripts can be accessed by the link: https://github.com/guandailu/nanopore_annotation.

## Notes

### Competing Interest Statement

The authors have declared no competing interest.

