## Supplemental Figures for "A comprehensive prediction of transcript isoforms in 19 chicken tissues by Oxford Nanopore long-read sequencing"

### Slide 1
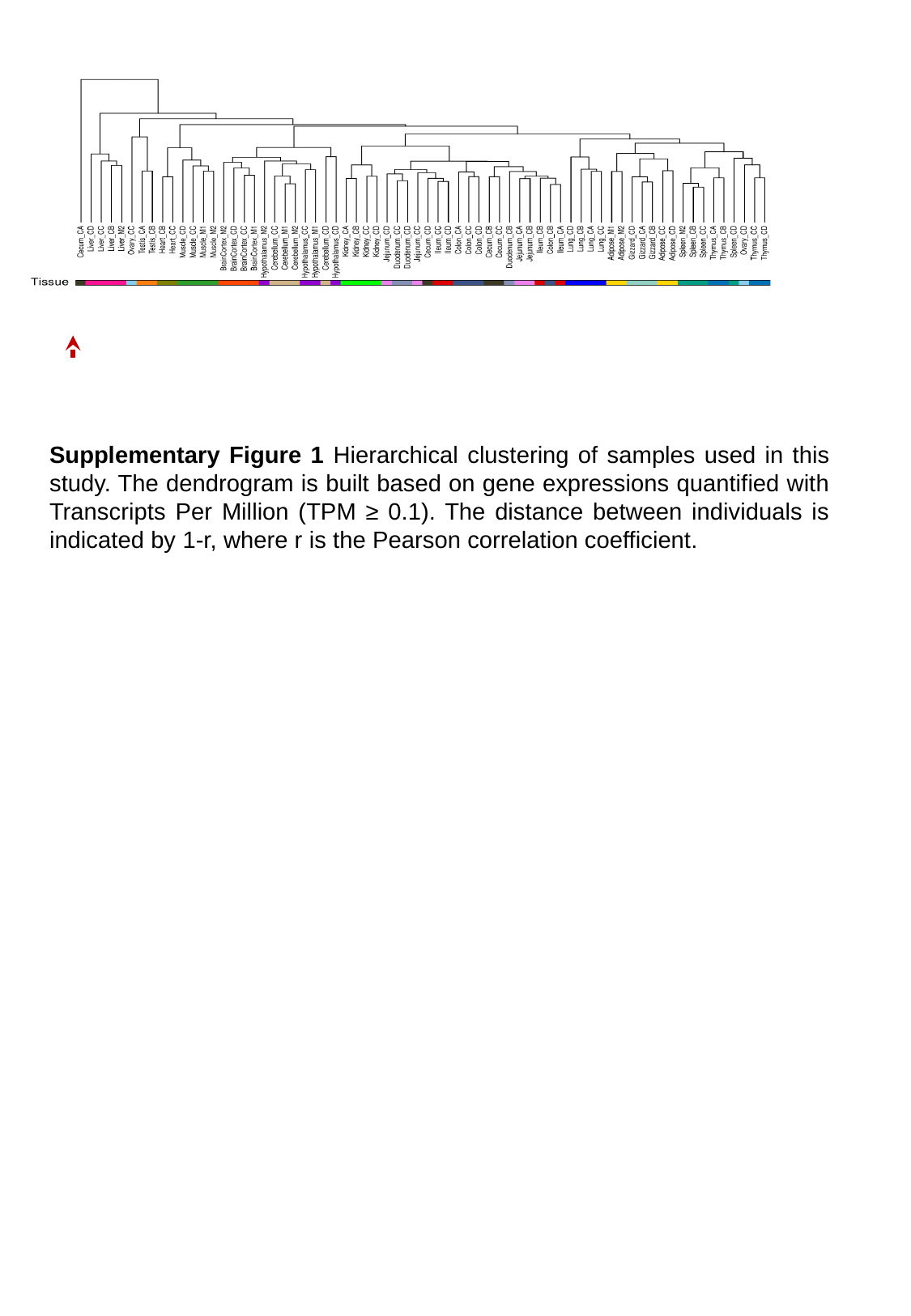

Supplementary Figure 1 Hierarchical clustering of samples used in this study. The dendrogram is built based on gene expressions quantified with Transcripts Per Million (TPM ≥ 0.1). The distance between individuals is indicated by 1-r, where r is the Pearson correlation coefficient.

### Slide 2
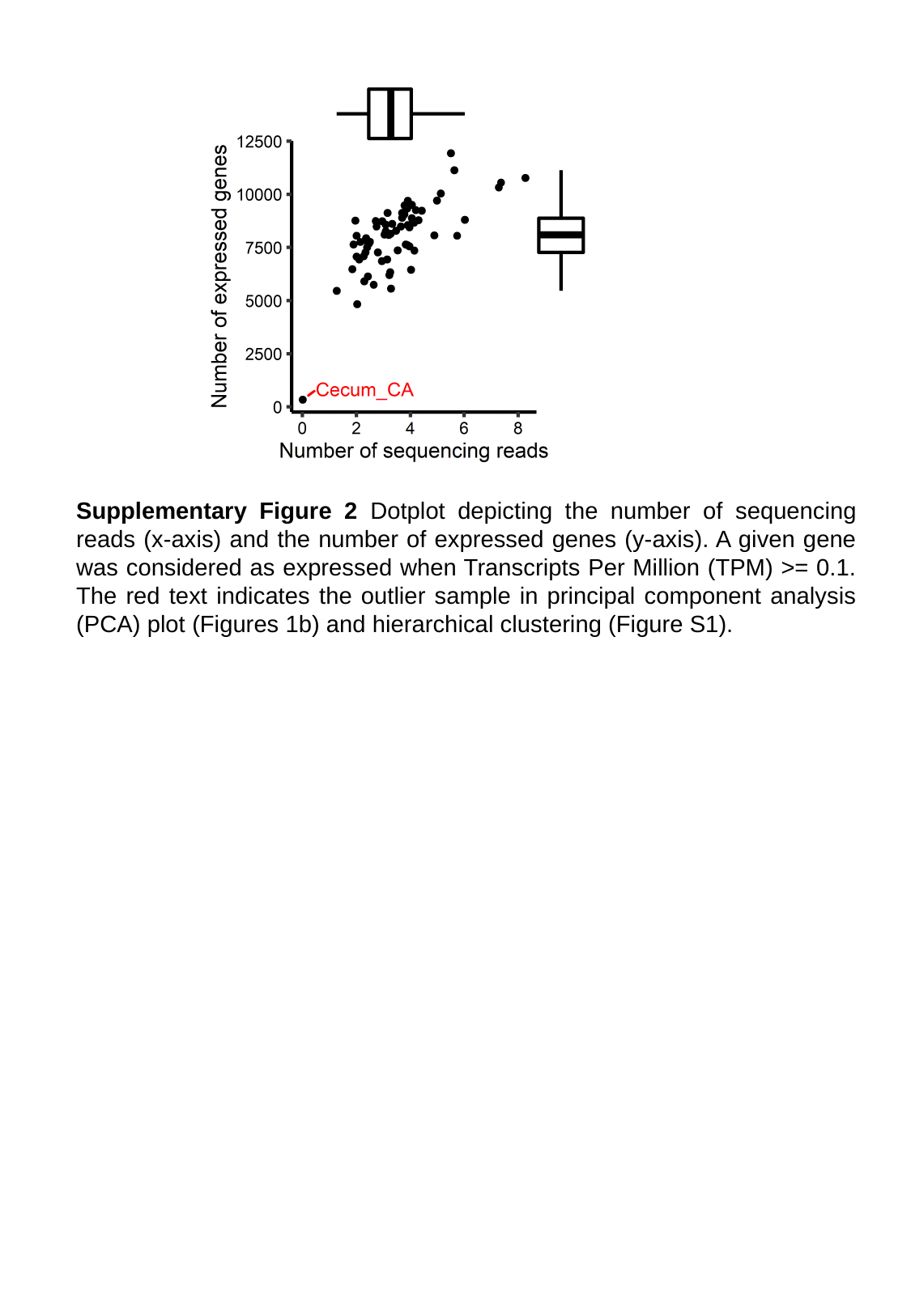

Supplementary Figure 2 Dotplot depicting the number of sequencing reads (x-axis) and the number of expressed genes (y-axis). A given gene was considered as expressed when Transcripts Per Million (TPM) >= 0.1. The red text indicates the outlier sample in principal component analysis (PCA) plot (Figures 1b) and hierarchical clustering (Figure S1).

### Slide 3
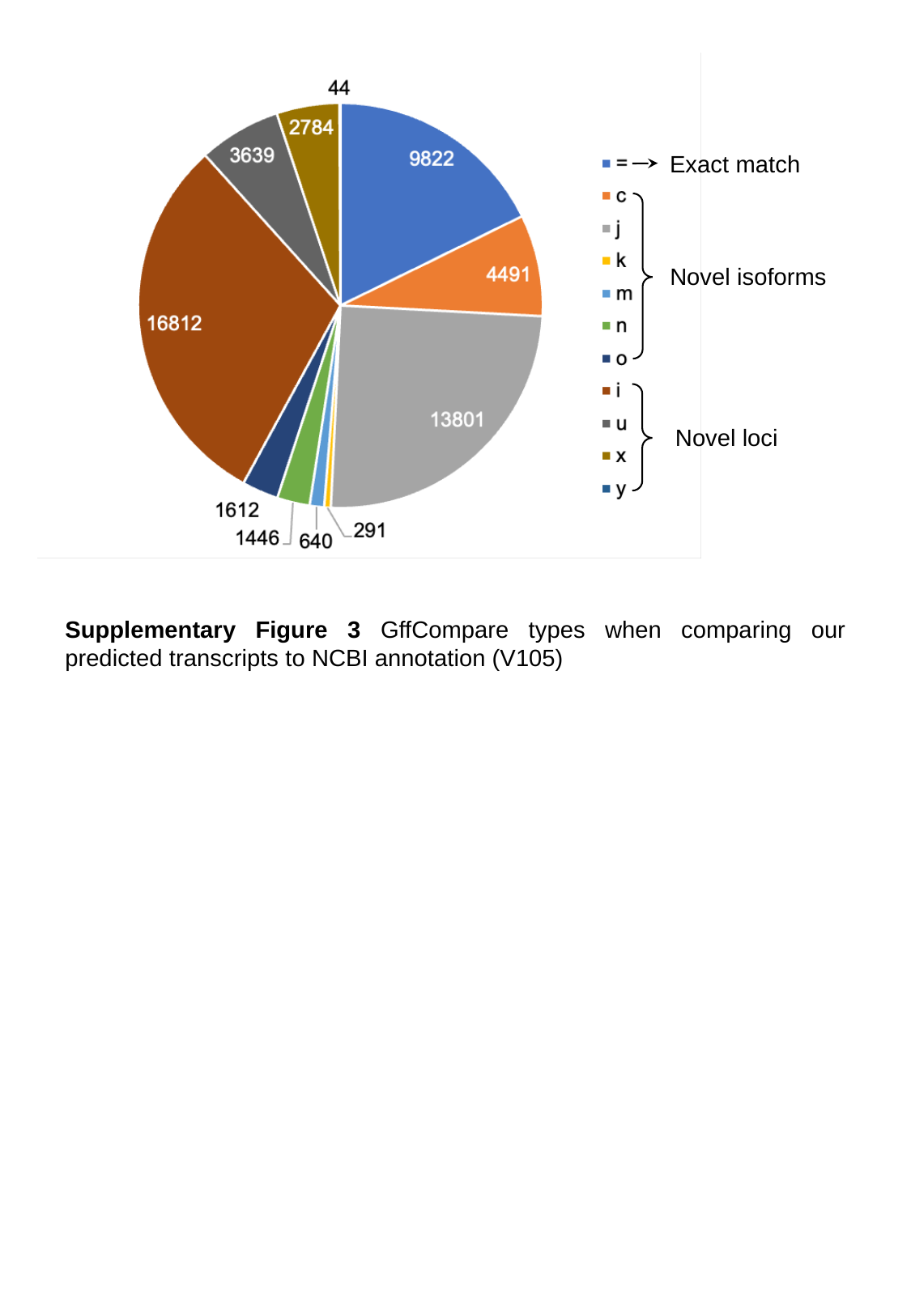

Exact match
Novel isoforms
Novel loci
Supplementary Figure 3 GffCompare types when comparing our predicted transcripts to NCBI annotation (V105)

### Slide 4
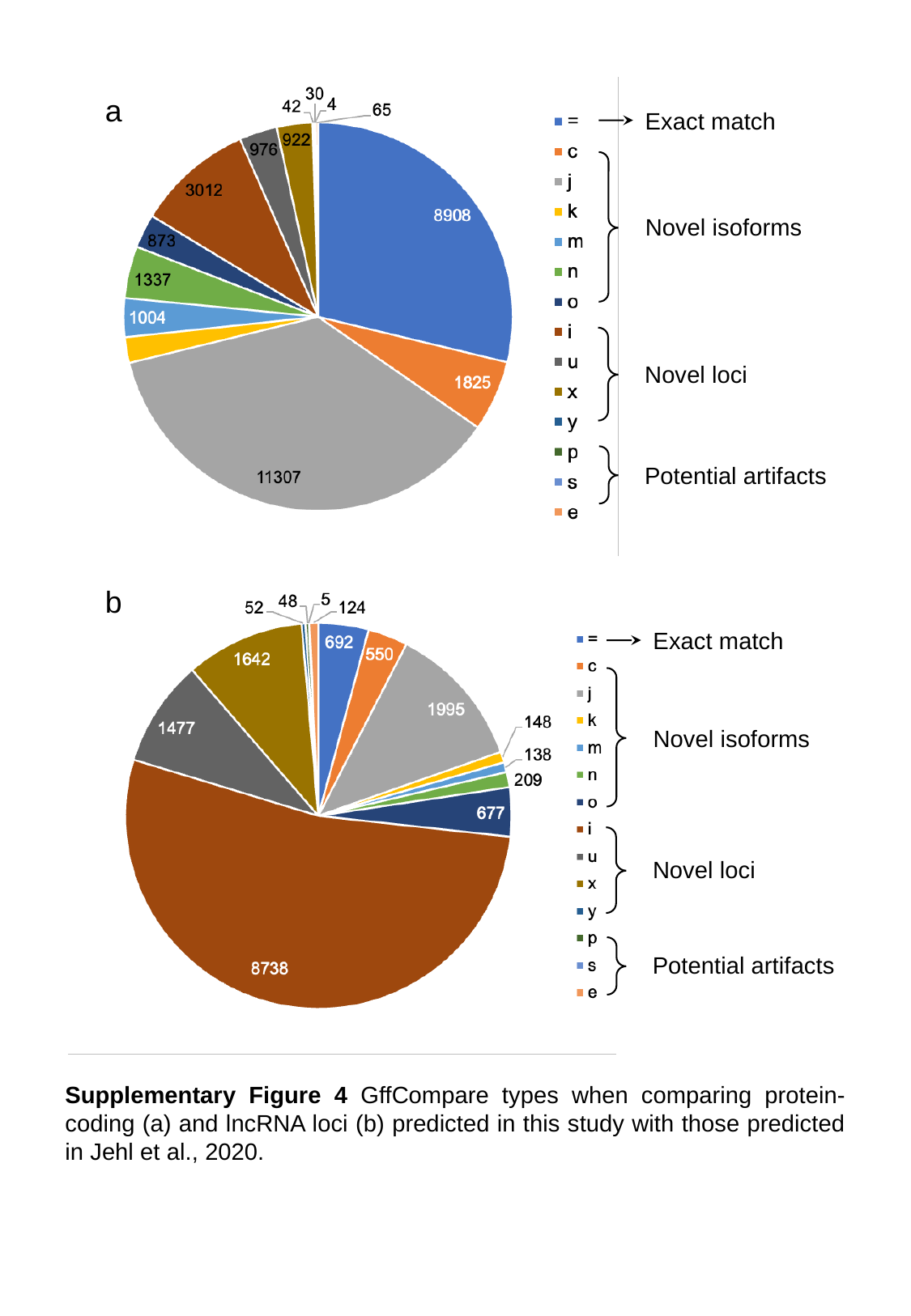

a
Exact match
Novel isoforms
Novel loci
Potential artifacts
b
Exact match
Novel isoforms
Novel loci
Potential artifacts
Supplementary Figure 4 GffCompare types when comparing protein-coding (a) and lncRNA loci (b) predicted in this study with those predicted in Jehl et al., 2020.

### Slide 5
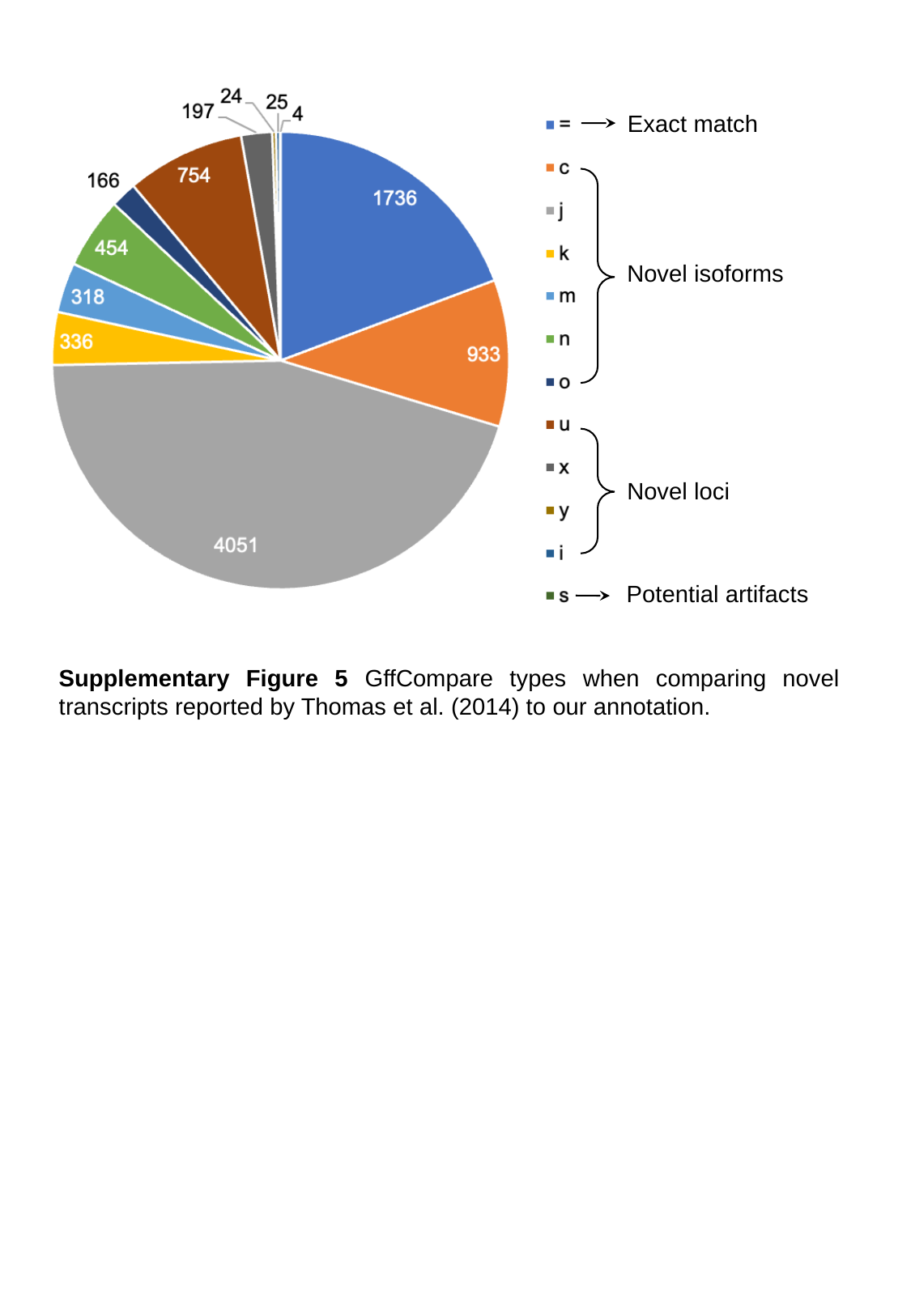

Exact match
Novel isoforms
Novel loci
Potential artifacts
Supplementary Figure 5 GffCompare types when comparing novel transcripts reported by Thomas et al. (2014) to our annotation.

### Slide 6
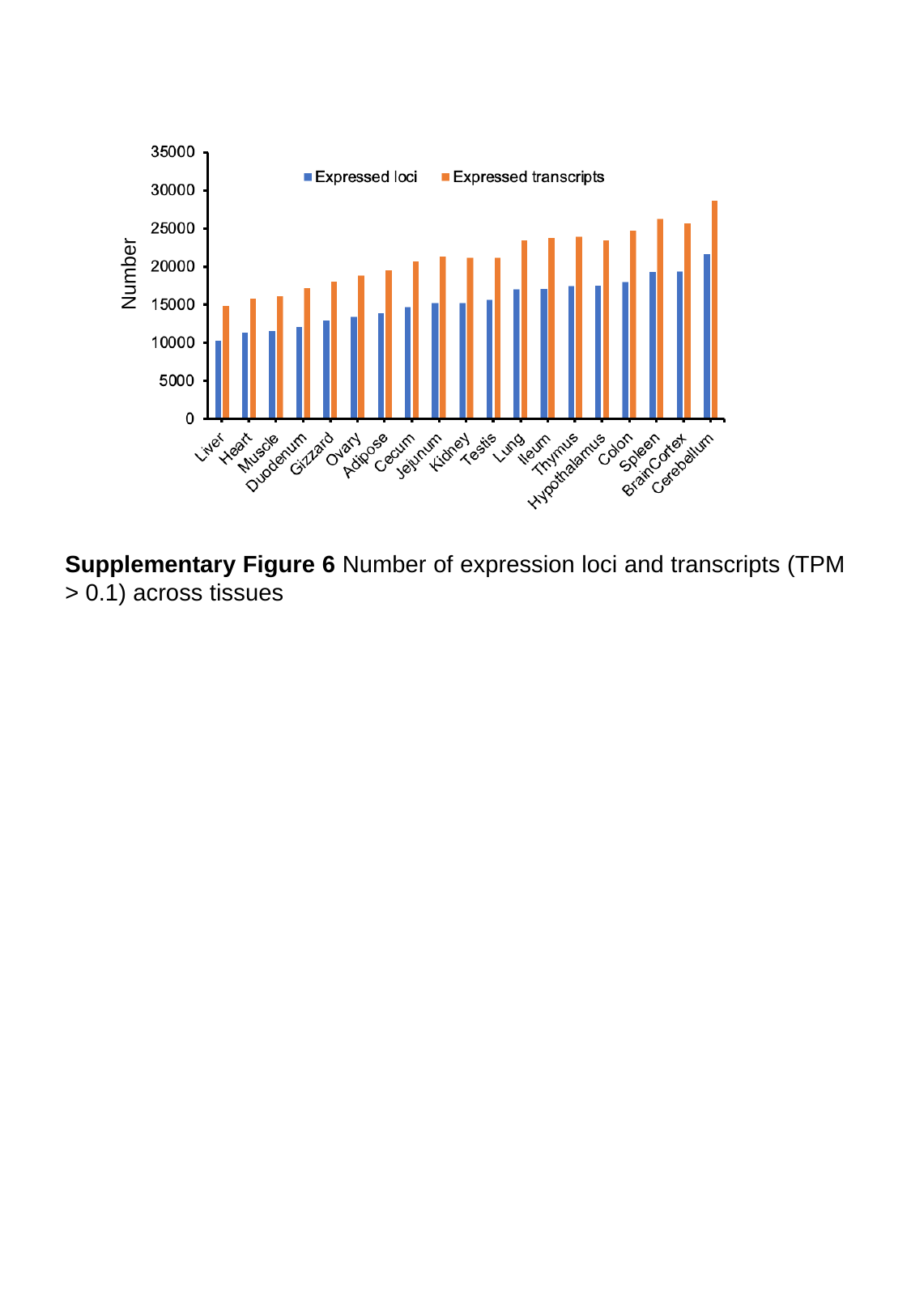

Number
Supplementary Figure 6 Number of expression loci and transcripts (TPM > 0.1) across tissues

### Slide 7
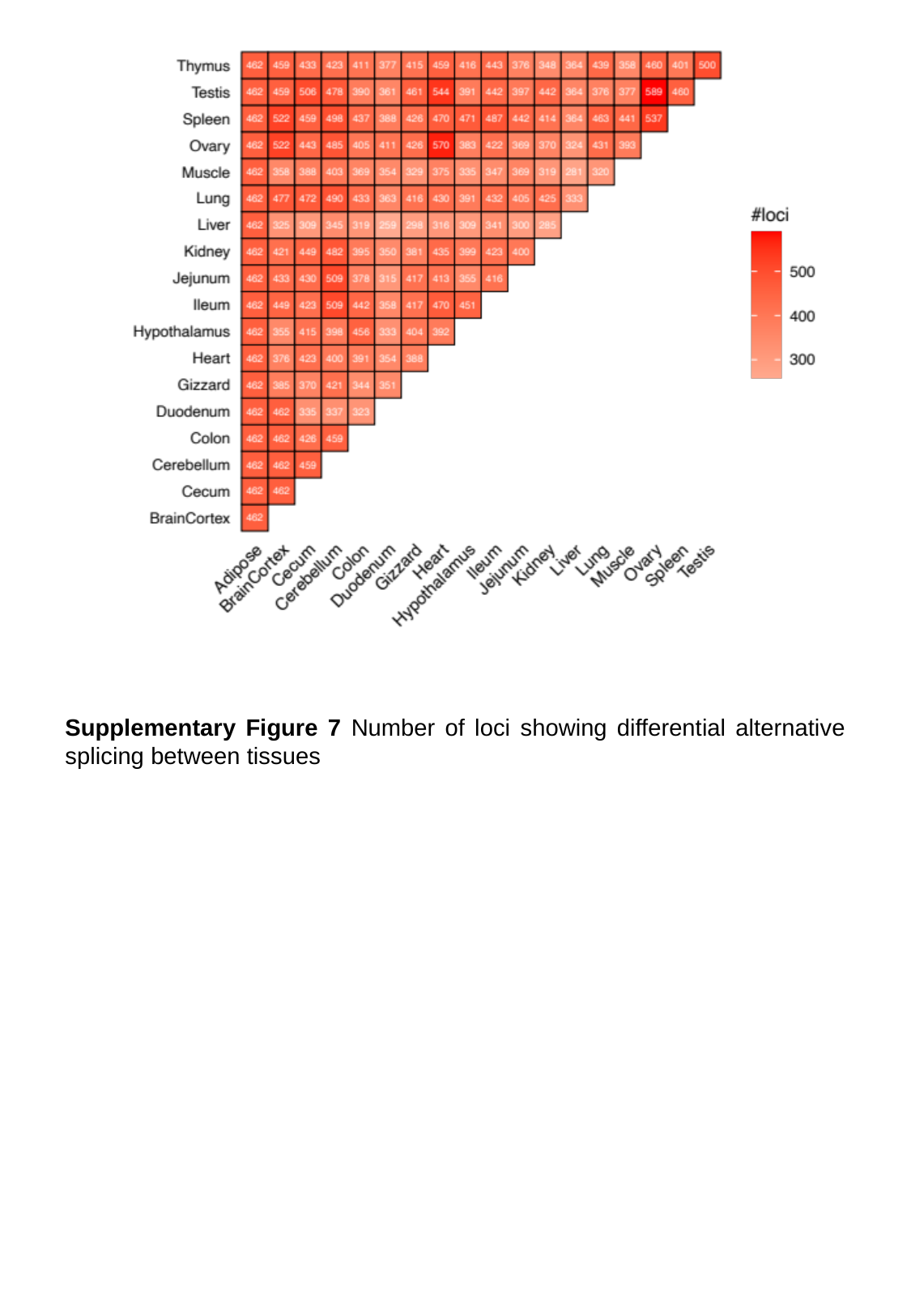

Supplementary Figure 7 Number of loci showing differential alternative splicing between tissues
